# Functional effects of TBC1 domain containing kinase deletion in immortalized B cells and plasma cells

**DOI:** 10.1101/2023.10.23.563612

**Authors:** Troy von Beck, Joshy Jacob

## Abstract

TBC1 domain containing kinase (TBCK) is a ubiquitous protein pseudokinase highly expressed in neurons and glial cells. Mutations in TBCK disrupt nervous system function and cause a characteristic syndrome of intellectual disability and hypotonia with various other comorbidities occurring in non-nervous tissues. Previous studies have shown a vacuolation defect present in B cells from individuals with TBCK mutations, but it is unclear if this affects plasma cell differentiation or humoral immunity. Recent research has revealed that TBCK is part of the FERRY complex involved in mRNA transport and is implicated in regulating mTORC1 signaling, autophagy, and cellular processes like cell proliferation and migration. Yet, the exact role of TBCK in these processes is still not fully understood. In this study, TBCK knockout cell lines were generated to investigate its impact on B cells and plasma cells. TBCK knockout plasma cells showed reduced immunoglobulin secretion after one week in culture, suggesting a possible defect in recycling cell components or energy usage. However, more experiments are needed to confirm this observation.

## Introduction

TBC1 domain containing kinase (TBCK) is ubiquitously expressed protein containing a conserved protein kinase, rhodanese, and eponymous Tre-2/Bub2/Cdc16 (TBC) domain. Despite its low tissue specificity, TBCK is roughly 10-20-fold more highly expressed in neurons and glial cells, a factor which underlies the severe intellectual disability and hypotonia in rare cases of bi-allelic loss-of-function mutations. While the most severe symptoms derive from neuronal dysfunction, it is clear other tissues are also affected from the apparent craniofacial dysmorphia, skeletal perturbations, hypothyroidism, dyslipidemia, and sporadic mentions of immunodeficiency^1-3^. A previous characterization of two pediatric individuals with homozygous TBCK loss-of-function mutations identified a severe vacuolation defect in the B-cell but not T-cell or myeloid lineages. Specifically, this vacuolation defect appeared in CD20^+^ B cells and CD138^lo^ immature plasma cells^4^. Currently it is unclear whether the absence of vacuolated mature plasma cells in TBCK syndrome patients indicates a block in plasma cell differentiation or alternatively, a rescue of the non-vacuolated phenotype incurred upon plasma cell differentiation. Further, it is unknown the extent to which TBCK loss-of-function mutations impact humoral immunity as the authors of that study did not report whether this defect was associated with an alteration in serum antibody concentrations or susceptibility to infection.

Recent molecular studies have identified that TBCK is a protein subunit of the FERRY complex responsible for linking mRNAs to early endosomes via Rab5 and thereby facilitating their intracellular transport. Further, this complex appears to selectively associate with a subset of mRNAs enriched for nuclear-encoded mitochondrial proteins and components of the nucleosome and endosomal system^5,6^. While the exact role of TBCK in this complex is largely undetermined, the protein appears to function in the regulation of several key cellular processes. First, TBCK promotes mTORC1 signaling and S6 phosphorylation in patient-derived lymphoblastoid cell lines and absence of TBCK results in increased autophagosome formation and impaired oligosaccharide degradation in patient-derived fibroblasts^2,3^. This modulation of mTORC1 signaling may be a downstream effect of epidermal growth factor (EGF) receptor signaling, as TBCK appears to link EGF receptor stimulation to ERK1/2 phosphorylation and attenuation of STAT3 phosphorylation^7^. This is further supported by observations of TBCK syndrome patient derived induced neuroprogenitor cells (iNPCs), which had an exacerbated loss of S6 phosphorylation following growth factor withdrawal^8^. Functionally, loss of TBCK in iNPCs or HEK293 cells produces deficits in cell proliferation, migration, and cell size, where the latter was linked to altered F-actin organization^8,9^. Finally, it should be noted that while TBCK contains a putative protein kinase domain, it is likely to be non-functional due to mutations in several highly conserved catalytic residues. Similarly, the putative Rab-GAP domain lacks a known Rab GTPase partner^9^.

Despite the clear significance of TBCK in cell homeostasis, it is unclear whether loss-of-function mutations have a profound impact on antibody production. In the following experiments we have attempted to define a B cell model of TBCK deficiency. CRISPR-Cas9 targeting the TBCK locus was used to generate TBCK knockout “Raji” B cell and “LP-1” plasma cell lines. Despite what has been described in studies of other cell lines, all TBCK knockout modified cell lines in our study had similar proliferation, survival, and sensitivity to serum withdrawal as their wild-type progenitors. However, modified LP-1 cells appeared to have reduced immunoglobulin secretion after 1 week in culture, perhaps indicating an inability to efficiently recycle cell components or use alternate energy sources in spent media. However, at the time of writing that observation is based on a single experiment performed in duplicate.

## Results

To investigate TBCK in B cell function we utilized CRISPR-Cas9 delivered by a lentiviral vector to target the TBCK locus using one of several guide RNAs (Figure 1) in either the Epstein-Barr virus transformed “Raji” B cell line or the multiple myeloma “LP-1” cell line. These two cell lines represent distinct phases in B cell differentiation, where Raji cells are akin to circulating naïve or memory B cells and the LP-1 are akin to plasma cells and secrete large amounts of IgG1 isotype immunoglobulin. For Raji cells, lentiviral transduction resulted in substantial reductions in TBCK expression for cells transfected with guide RNAs KO2, KO4, KO5, or KO6 (Figure 2A-B). For LP-1 cells, knockdown of TBCK expression after lentiviral transduction was initially unimpressive and was only modestly reduced in cells receiving either the KO4 or KO6 guide RNAs. To improve the knockdown of TBCK expression in LP-1 cells, clones of each LP-1 TBCK variant were produced by limiting dilution and expansion from a single cell. Several of these clones possessed satisfactory knockdown of TBCK expression and were utilized in later analyses (Figure 2C-D).

**Figure 1.**
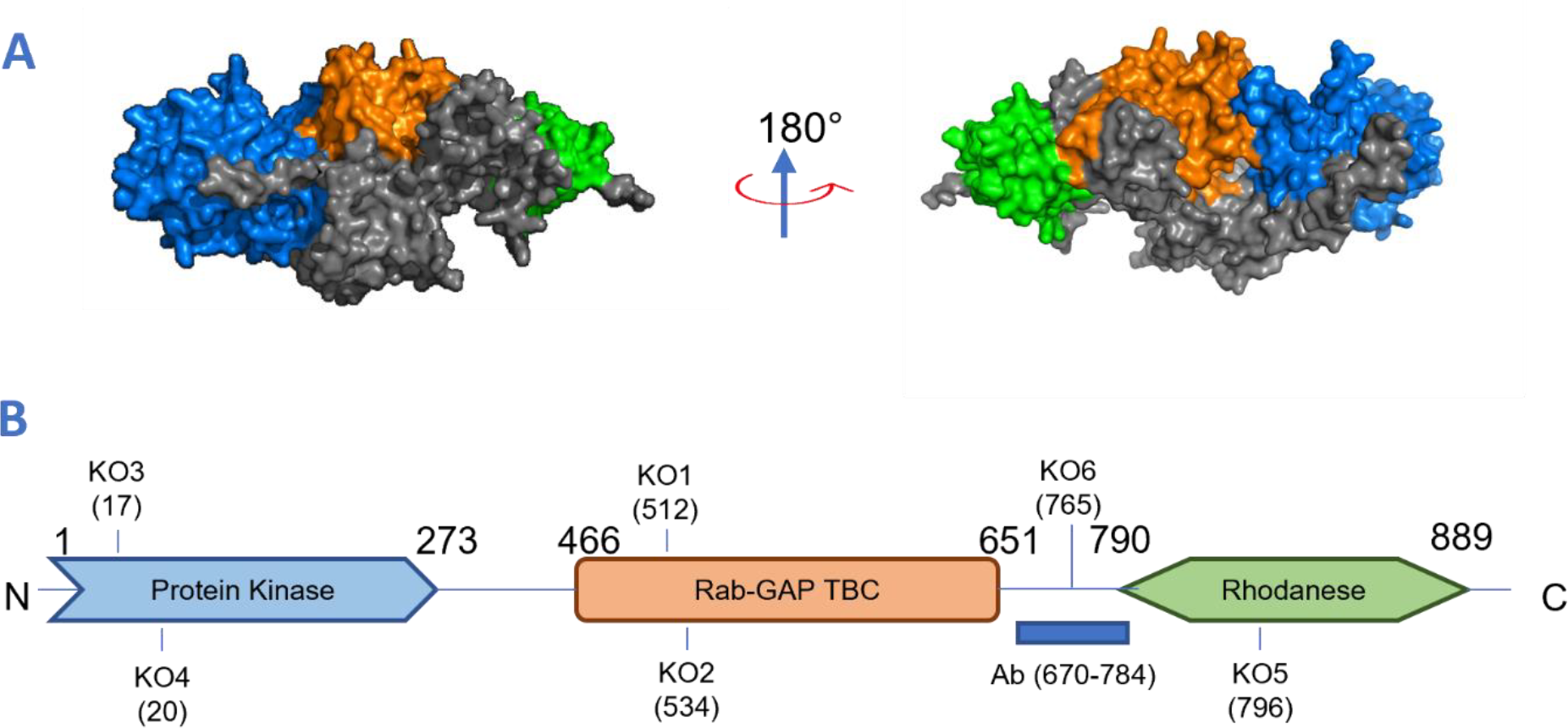
3D structure of TBCK and location of CRISPR modified variants characterized in this study. (A) 3D structure of TBCK as predicted by AlphaFold (Q8TEA7). Protein kinase (blue), Rab-GAP TBC (orange), and rhodanese (green) domains are depicted. (B) Schematic representation of the long isoform of TBCK with call outs for the residues targeted by each CRISPR construct and the α-TBCK antibody.

**Figure 2.**
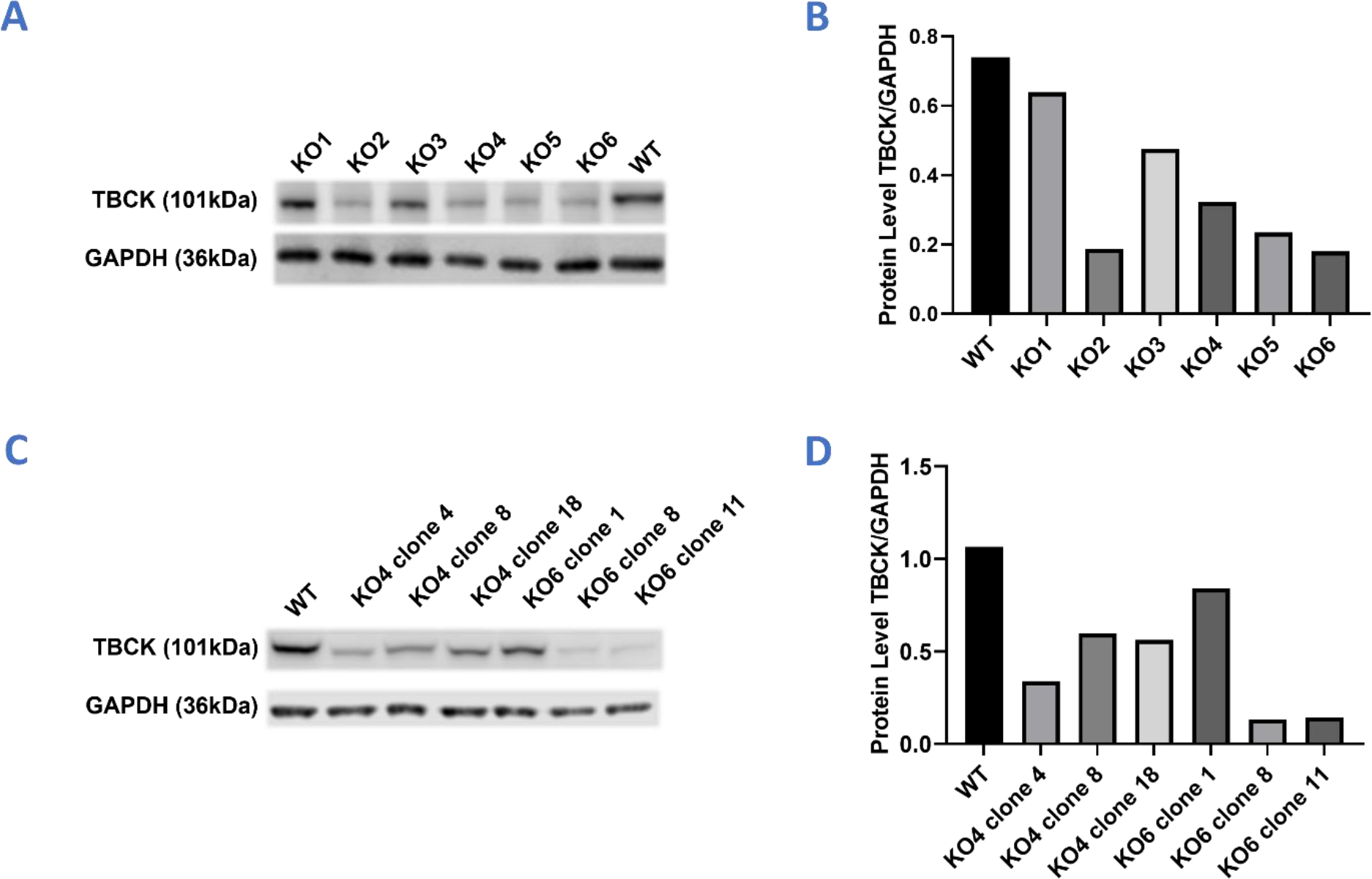
Characterization of TBCK expression in CRISPR edited B cell lines. (A-B) Western blot of TBCK protein expression in CRISPR modified Raji cell lines. (C-D) Western blot of TBCK protein expression in CRISPR modified LP-1 cell lines. Densitometry measurements were taken in ImageJ version 1.53f51.

We first examined these cells by flow cytometry to determine whether the effects of TBCK knockdown observed in previous patient-derived or immortalized-cell models could be reproduced in Raji or LP-1 cells. For this experiment, Raji or LP-1 cells were grown for 24 hours in media containing either 10% FBS or 0% FBS to simulate conditions with and without EGFR signaling. Curiously, neither Raji (Figure 3A-C, Figure S1A) nor LP-1 (Figure 3D-G, Figure S1B) cells displayed any sensitivity to the loss of TBCK either under normal or no-serum stress conditions. All modified cell lines had similar degrees of viability, cell size, and phosphorylation of S6. Further, modified cell lines had similar side-scatter profiles, implying there was no increase in cellular granularity, and therefore no accumulation of autophagosomes in the TBCK knockout cells. Consistent with previous reports, both cell lines exhibited drops in S6 phosphorylation following serum withdrawal, and this effect was more pronounced in LP-1 cells, perhaps indicating a greater reliance on EGF for homeostasis.

**Figure 3.**
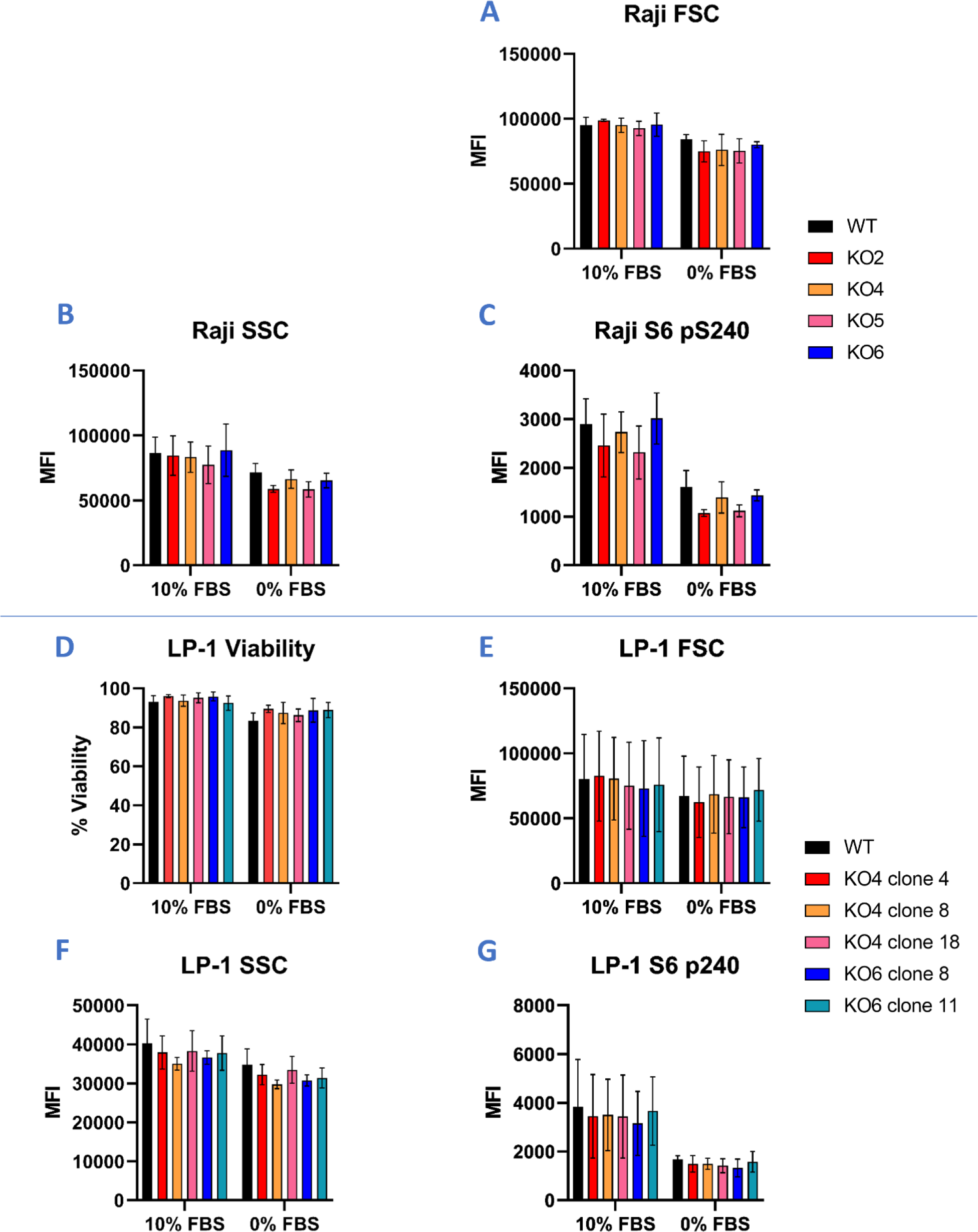
Loss of TBCK does not alter S6 phosphorylation in Raji or LP-1 cells. (A-C) Raji cell size (FSC), complexity (SSC), and S6 S240 phosphorylation after 24 hours in either 10% or 0% FBS containing DMEM. (D-G) LP-1 cell viability, size (FSC), complexity (SSC), and S6 S240 phosphorylation after 24 hours in either 10% or 0% FBS containing DMEM. Graphs represent the mean and standard deviations from three independent experiments.

Given that neither cell line appeared meaningfully impacted by the loss of TBCK in terms of cellular granularity, size, mTOR activity, or sensitivity to EGF withdrawal, we further characterized the functional effects of TBCK loss on cellular proliferation and immunoglobulin secretion using the LP-1 cell line. LP-1 cells were again grown in conditions of either high (10%) or low (1%) serum abundance to mimic normal and stress-inducing conditions but were also grown in either atmospheric oxygen or hypoxic (2.5% O_2_) conditions to mimic the bone marrow environment where long-lived plasma cells reside. Like the flow cytometric analysis, LP-1 cells proliferated to similar extents over the course of 12 days under all tested conditions as measured by an MTT formazan formation assay (Figure 4A). Interestingly 1% FBS conditions were growth limiting under normoxic, but not hypoxic conditions, indicating that oxygen availability became the rate limiting factor in cellular metabolism.

**Figure 4.**
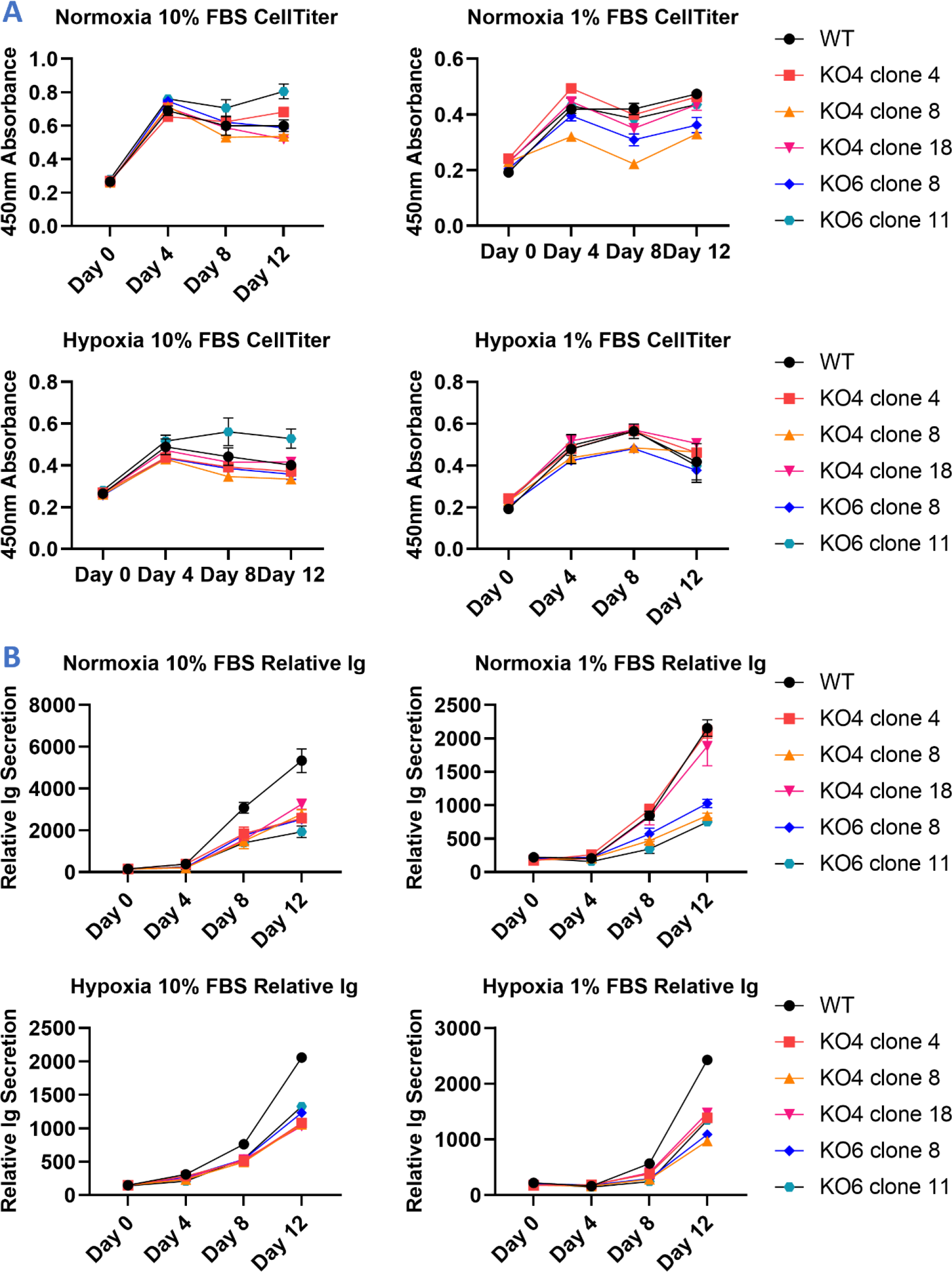
Survival and Immunoglobulin secretion of WT and TBCK KO LP-1 cells. (A) LP-1 cells were grown in 10% or 1% FBS and normoxia or hypoxia conditions for 12 days. On days 0, 4, 8, and 12, cell supernatants were collected and metabolic activity measured by MTT assay. For hypoxic conditions, the MTT assay was also performed under hypoxia. (B) Supernatants from LP-1 cell cultures were analyzed by quantitative ELISA to determine the IgG1 concentrations. IgG1 titers were normalized to the metabolic activity presented in panel A to account for potential differences in cell number at seeding or due to differential proliferation. Graphs are representative of 1 experiment performed in duplicate.

Despite the absence of TBCK mediated effects on cellular proliferation and metabolism, loss of TBCK appeared to cause a reduction in antibody production. Up to day 4, the production of immunoglobulin normalized to MTT metabolic activity remained consistent between WT and TBCK knockout cell lines (Figure 4B). However, on days 8 and 12, WT cells began to show a substantial increase in the production of immunoglobulin relative to their metabolic activity. This effect was consistent across all culture conditions except for normoxia with 1% FBS, where KO4 clone 4 and KO4 clone 18 achieved comparable immunoglobulin production. If this effect is reproducible, it may represent a distinct metabolic requirement for TBCK to achieve effective antibody secretion under nutrient limiting conditions.

## Discussion

Here we have established TBCK deficient cell lines to study the role of this protein at two distinct phases of B cell differentiation using Raji and LP-1 cells. Surprisingly, the results observed for the derivative cell lines conflict with both patient-derived samples and previous studies of modified immortalized cell lines representing neuronal, epithelial, and even lymphoblastoid type cells^2-4,8,9^. The absence of increased granularity in either the Raji or LP-1 cells suggests that some condition for TBCK deficiency driven autophagosome formation is not met. One explanation may be that both Raji and LP-1 cells have acquired mutations which make them insensitive to TBCK deficiency, however the absence of such mutations in other immortalized cell lines would suggest otherwise. More likely, we hypothesize that the vacuolation phenotype represents a transient stage in B cell differentiation, potentially representing some stage of B cell activation and terminating at the point of plasma cell differentiation.

Curiously, while LP-1 cells appear insensitive to TBCK deletion using previously described metrics, they do appear to require TBCK for optimal antibody secretion during longer incubations in cell culture. As the LP-1 deficient cells survived and proliferated to similar extents as their wild-type counterparts, it can be assumed the decrease in antibody production reflects the efficiency of antibody production rather than differences in cell population. Notably the reduction in relative antibody secretion was not apparent until day 8 in cell culture and continued to increase on day 12 in a largely uniform manner across culture conditions. This likely indicates that the expression of TBCK better allows LP-1 cells to maintain homeostasis and support antibody production under nutrient limiting conditions for key metabolites in RPMI based cell media. In support of this, we note that the discrepancy in antibody production appears after day 4, which also coincides with the termination of log-phase growth as determined by the stable MTT assay results from days 4-12.

While we have performed some preliminary characterization of TBCK function B cell lines, there are several limitations to the current study. Most notably, the observations presented in figure 4 are derived from a single experiment and therefore should not be considered definitive. Further, the knockout cell lines generated retained some residual TBCK production, ranging between roughly 10-50% depending on the cell line, which could mask specific phenotypes. Finally, the LP-1 knockout cell lines are monoclonal derivatives of the polyclonal wild-type precursors, and we cannot rule out that the observed differences in antibody production are due to the random selection of “low” antibody secreting clones from the parent population.

## Methods

### Cell lines and culture conditions

Low passage number Raji cells (Cat#: CCL-86, ATCC) were kindly provided by Dr. Chris Scharer. Low passage number LP-1 cells were kindly provided by Dr. Larry Boise. Raji and LP-1 cells were maintained in RPMI + 10% FBS, 1mM L-glutamine, and 1X penicillin-streptomycin. All cells were grown at 37°C and 5% CO_2_ unless otherwise stated.

### Design and synthesis of CRISPR lentiviral plasmids

Guide RNAs for gene knockouts targeting the 3 major domains of TBCK were chosen using the CHOPCHOP web tool for CRISPR/Cas9^10^. The following target sequences were each cloned into the doxycycline inducible LentiCRISPR v2 plasmid, TCLV2 (Cat#:87360, Addgene): KO1, 5’-TCATCGTACTGATGACAGCGAGG-3’; KO2, 5’-TAGGCGTGTATTAAAAGCCTGGG-3’; KO3, 5’-AACATCATGTGGCAGAGCCGAGG-3’; KO4, 5’-TTGCTTCCACAAACATCATGTGG-3’; KO5, 5’-TATTCCGGATGTCAACCACCAGG-3’; KO6, 5’-ATCACCACGGATTTCAGCAGAGG-3’.

### Production of CRISPR lentivirus

Lentivirus was produced in T75 flasks containing 4 x 10^6^ HEK293T cells seeded on the previous day. Cell media was replaced with fresh DMEM containing 5% FBS 1 hour prior to transfection. TCLV2 KO plasmids and the lentiGuide-Thy1.1 base editor gRNA plasmid were complexed with the psPAX2 and pMD2 plasmids for 3^rd^ generation lentivirus production in a 4:3:1 ratio. HEK293T cells were transfected using the Transporter 5 transfection reagent (Cat#: 26008, Polysciences) according to the manufacturer’s recommendations. Cells were then incubated at 37°C and 5% CO_2_ for 72 hours prior to collection of lentivirus containing supernatants. Supernatants were clarified at 3,000 x *g* for 10 minutes and filtered through a 0.45µm pore PVDF filter. Aliquots were frozen at -80°C until use.

### Generation of TBCK modified cell lines

Raji or LP-1 cells growing in RPMI + 10% FBS were seeded at a density of 3 x 10^5^ cells/mL and supplemented with 6µg/mL of polybrene. Lentiviral containing supernatants were then added, and the cultures were incubated for 72 hours. After 72 hours, cells infected with TCLV2 vector DNA were supplemented with 2µg/mL of puromycin to kill non-transfected cells. For TCLV2 infected cells, Cas9 expression was induced by the addition of doxycycline to a concentration of 5µg/mL. After 72 hours of doxycycline induction, GFP positive cells were sorted by flow cytometry.

### Western Blot

1 x 10^6^ Raji or LP-1 were pelleted and resuspended in 100µL of ice cold RIPA buffer containing 1X Halt protease inhibitors (Cat#: 78430, ThermoFisher Scientific) and incubated on ice for 30 minutes. Cell lysates were clarified at 2,000 x rcf for 3 minutes and supernatants collected and stored at -20° C until analysis. Denaturation of cell lysates in RIPA buffer was achieved by addition of NuPAGE LDS sample buffer and 5% β-mercaptoethanol with heating at 95°C for 5 minutes. Proteins were then separated by loading 15µL of denatured sample on a NuPAGE 4-12% Bis-Tris gel and transferred to a nitrocellulose membrane according to the manufacturer’s recommendations. Membranes were blocked with TBST + 5% BSA for 1 hour at room temperature and then incubated overnight at 4°C with primary antibody diluted 1:1,000 in TBST + 3% BSA. Unbound primary was removed by 3 washes with TBST + 3% BSA. Membranes were then incubated for 1 hour at room temperature with secondary antibody diluted 1:10,000 in TBST + 3% BSA. Unbound secondary was removed by 3 washes with TBST. Western blots were developed using chemiluminescence with the SuperSignal West Femto Substrate (Cat#: 34095, ThermoFisher) for detection of HRP conjugated secondary antibodies or using fluorescence in the 800nm channel for detection of IRDye 800CW conjugated secondary antibodies. Antibodies: α-TBCK (Cat#: HPA039951, Sigma-Aldrich), α-GAPDH (Cat#: 649201, BioLegend), α-rabbit Ig HRP (Cat#: 4030-05, SouthernBiotech), α-mouse Ig IRDye800CW (Cat#: 926-32210, LI-COR)

### Serum deprivation assay

200,000 Raji or LP-1 cells grown in RPMI + 10% FBS for at least 24 hours were pelleted and resuspended in either 1 mL of RPMI + 10% FBS or RPMI without FBS supplementation. Cells were incubated overnight at 37°C and 5% CO_2_. 24 hours later, cells were collected and pelleted by centrifugation at 300 x *g* for 5 minutes. Because Raji and LP-1 cells grown without FBS become adherent, all wells were further washed with 1X PBS + 10 mM EDTA and gentle pipetting to collect the adherent cells. Cell pellets were immediately processed for flow cytometric analysis.

### Flow Cytometry Quantification of RPS6 phosphorylation

Raji or LP-1 cell pellets were resuspended and stained with LIVE/DEAD near-IR (Cat#: L10119, ThermoFisher Scientific) viability stain prior to fixation in 1mL of BD Cytofix Fixation Buffer (Cat#: 554655, BD Biosciences) for 15 minutes at 37°C. Fixed cells were pelleted and then washed twice with BD Phosflow Perm/Wash Buffer I (Cat#: 557885, BD Biosciences). Cells were stained in a 20µL volume containing 4µL of Alexa Fluor 647 Mouse anti-S6 (pS240) (Cat#: 560432, BD Biosciences) and 16µL of perm/wash buffer. Cells were stained for 1 hour at room temperature, washed once with perm/wash buffer, and resuspended in 1X PBS + 1% FBS. Flow cytometric readings were captured on a BD LSR II flow cytometer using the APC channel to detect AF547 staining. Analyses were performed in FlowJo v10.7.1.

### MTT metabolic activity and Ig secretion analysis

LP-1 cells were split 1 day prior to seeding to ensure uniform culture conditions. On the day of, cells were pelleted and resuspended in fresh media and seeded at 1 x 10^5^ cells/mL in 96-well plates with a 100µL culture volume. For culture conditions the media contained either 10% or 1% FBS and was maintained in either 5% CO_2_ with atmospheric oxygen or in 5% CO_2_ with only 2.5% oxygen using a hypoxia incubator. Immediately after plating and on days 4, 8 and 12, 10µL of cell supernatants were collected for analysis of Ig secretion by ELISA. Similarly, the number of cells was quantitated on day 0, 4, 8, and 12 by MTT formazan formation assay by adding 20µL of CellTiter 96 Aqueous One solution (Promega, Catalog#: PAG3580), incubating for 1 hour, and reading the absorbance at 490nm. The concentration of Ig in serum supernatants was measured using a standard sandwich ELISA with anti-human Ig primary (Cat#: 2048-01, Southern Biotech) and secondary antibodies (Cat#: 2045-05, Southern Biotech) using a known concentration control (Cat#: 0150-01, Southern Biotech).

---

**Supplemental Figure 1.**
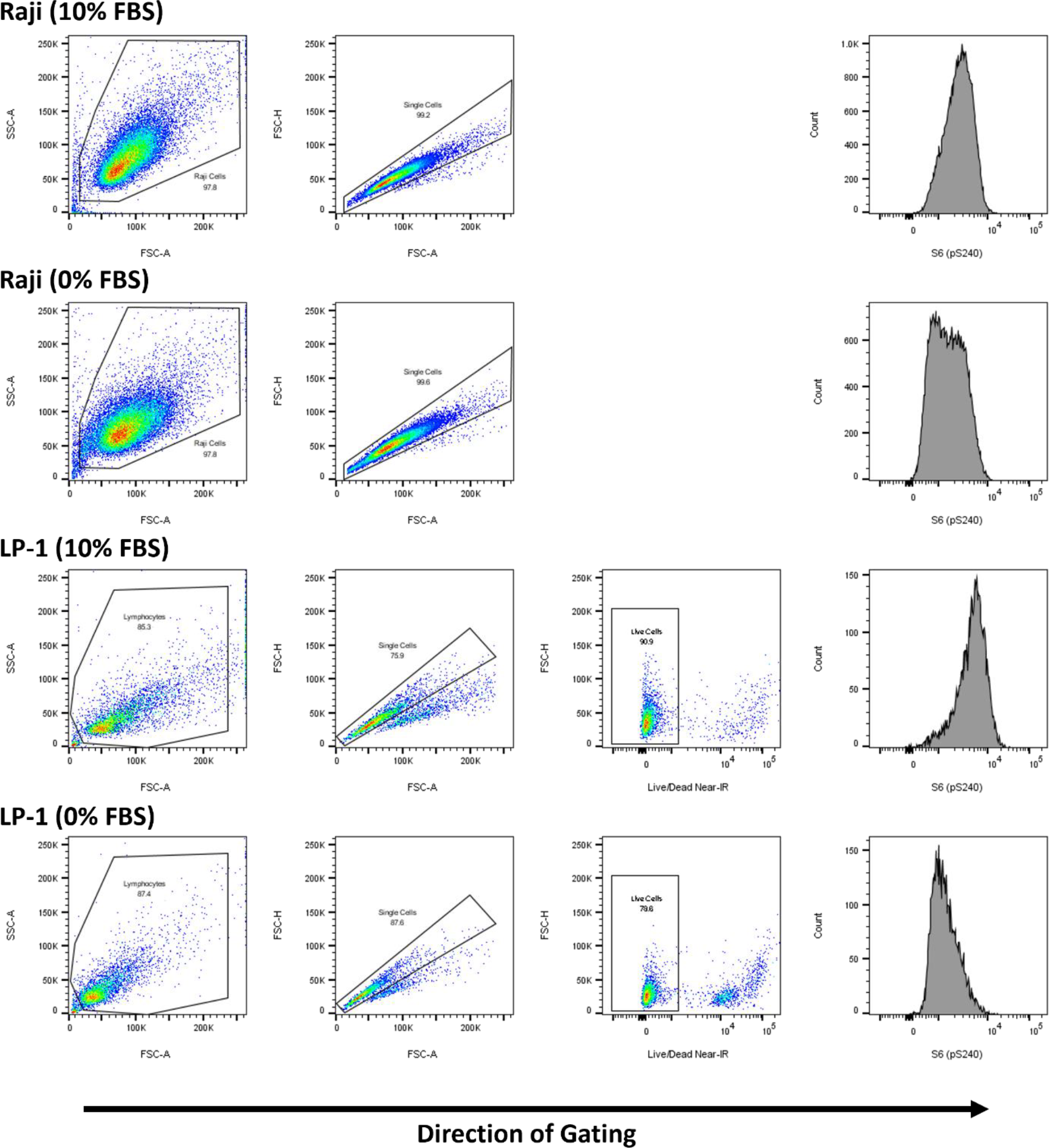
Representative flow cytometry plots for WT Raji and WT LP-1 cells after 24 hours in 10% and 0% FBS culture conditions. Viability was not recorded during Raji cell analysis.

## Notes

### Competing Interest Statement

The authors have declared no competing interest.

